# T-bet fate mapping identifies a novel ILC1-ILC2 subset *in vivo*

**DOI:** 10.1101/2020.08.21.261073

**Authors:** J-H Schroeder, N Garrido-Mesa, T Zabinski, AL Gallagher, L Campbell, LB Roberts, E Stolarczyk, G Beattie, JW Lo, A Iseppon, C Moreira Heliodoro, R Reis, RG Jenner, P Lavender, JK Howard, RK Grencis, H Helmby, J F Neves, GM Lord

## Abstract

Innate lymphoid cells (ILC) play a critical role in regulating immune responses at mucosal surfaces. Various subsets exist resembling T cell lineages defined by the expression of specific transcription factors. Thus, T-bet is expressed in ILC1 and Th1 cells. In order to further understand the functional roles of T-bet in ILC, we generated a fate-mapping mouse model that permanently marks cells and their progeny that are expressing, or have ever expressed T-bet. Here we have identified and characterised a novel ILC with characteristics of ILC1 and ILC2 that are “fate-mapped” for T-bet expression and arise early in neonatal life prior to establishment of a mature microbiome. These ILC1-ILC2 cells are critically dependent on T-bet and are able to express type 1 and type 2 cytokines at steady state, but not in the context of inflammation. These findings refine our understanding of ILC lineage regulation and stability and have important implications for the understanding of ILC biology at mucosal surfaces.

**SUMMARY:** Innate lymphoid cells (ILC) play a critical role in regulating immune responses at mucosal surfaces. Three distinct ILC groups have been described according to expression of subset defining transcription factors and other markers. In this study we characterize a novel ILC subset with characteristics of group 1 and group 2 ILC *in vivo*.

## INTRODUCTION

More than 30 years ago Mosmann and Coffman provided a fundamental framework for CD4^+^ T cell biology (Mosmann and Coffman, 1989). It has now been established that the immune response of helper T cells can be directed to either intracellular pathogens, extracellular helminths or small extracellular pathogens (e.g. fungi or bacteria). These diverse immune responses are driven by so called type 1, 2 or 17 immune responses respectively. All of these immune responses are characterized by specific sets of cytokines. In recent years, innate lymphoid cells (ILC) have been identified as a group of lymphocytes with analogous functions to CD4^+^ T cells (reviewed by McKenzie *et al.*, 2014). Group 1, 2 and 3 ILC (ILC1, ILC2 and ILC3) have been shown to drive a type 1, 2 or 17 immune response, respectively. Although CD4^+^ T cells and ILC have overlapping functions, there are fundamental differences. Unlike T cells, ILC do not express an antigen-specific receptor and are reliant on other signals and cells to direct their activation. These cells include antigen presenting cells, the epithelium including tuft cells, T cells and neuronal cells. Upon tissue damage the epithelium is the main provider of the ILC2-activating cytokines IL-33, IL-25 and TSLP, which together with the ILC2-activating mediator TL-1A can also be secreted by various APCs (reviewed by McKenzie *et al.*, 2014, Yu *et al.*, 2014). CD4^+^ T cells can also interact with ILC2 via the TCR-MHC class II axis resulting in the enhanced proliferation of ILC2 driven by IL-2 secreted by the interacting helper T cells (Oliphant *et al.*, 2014). Furthermore, IL-25 production by tuft cells can be triggered by succinate that appears in the gut lumen in case of microbial disbalance (Macias-Ceja *et al.*, 2019). Several neurotransmitters have also been suggested to promote the ILC response (reviewed by Vivier *et al.*, 2018, Klose *et al.*, 2020).

The presence of specific cytokines during T cell activation in secondary lymphoid organs is an important factor in polarizing helper T cells to Th1, Th2, Th17, Th9, Tfh or regulatory T cell subsets. However, these polarizations are reversible depending on the cytokines CD4^+^ T cells are being exposed to later in the tissue. This cytokine-driven phenomenon defined as plasticity has also been shown for ILC. As such, it has been observed that NKp46^+^ ILC3 can differentiate to ILC1 and vice versa (reviewed in Klose *et al.*, 2020), and likewise ILC2 have shown plasticity to ILC1 and ILC3 (Bal *et al.*, 2016, Silver *et al.*, 2016, Ohne *et al.* 2016, Lim *et al.*, 2016, Huang *et al.*, 2015 and Zhang *et al.*, 2018, Golebski *et al.*, 2019). In humans and mice, IL-12 and IL-18 appear to be the most potent factors to induce T-bet and IFN-γ in ILC2 (Bal *et al.*, 2016, Silver *et al.*, 2016, Ohne *et al.* 2016, Lim *et al.*, 2016). Some groups have also found that IL-4 derived from eosinophils may be instrumental to convert an ILC1 to an ILC2 (Bal *et al.*, 2016). This suggestion is interesting because ILC being predominantly resident in various mucosal organs, are likely to be in close contact with gut-resident eosinophils. It is known from studies on helper T cells that IL-4 is an important inducer of GATA3 expression (Bredo *et al.*, 2015).

It has been shown that plasticity in T cells is not always absolute. CD4^+^ T cells co-expressing T-bet and RORγt or T-bet and GATA3 expression have been characterized (Jenner et al 2009, Affinass *et al.*, 2018, Peine *et al.*, 2013). NKp46^+^ ILC3 are known to express RORγt and T-bet, but it is not known whether ILC exist that co-express GATA3 and T-bet. The existence of an ILC1-ILC2 has been predicted in RNAseq analyses using gut tissue and also from *in vitro* culture systems in which ILC were generated from FACS sorted human and murine ILC progenitors (Gury Ben-Ari *et al.*, 2016, Lim *et al.*, 2017, Xu *et al.*, 2019). In this study we identify a heterogeneous intestinal lamina propria ILC1-ILC2 subset that shares characteristic expression of surface markers and transcription factors as well as functional properties usually associated with ILC2 and ILC1. These ILC1-ILC2 cells are characterised by a distinct gene expression profile and are dependent on T-bet for their development and maintenance. ILC1-ILC2 cells arise early in neonatal life before full establishment of the microbiota. Although these cells express both type 1 and type 2 cytokines at steady state, they are dispensable for anti-helminth responses *in vivo* and appear to be less functional during colitis. The existence of ILC1-ILC2 highlights an important novel aspect of ILC biology that should be taken into consideration in the interpretation of studies into ILC subsets.

## RESULTS AND DISCUSSION

### Identification of T-bet-dependent NKp46-positive intestinal ILC2

Throughout this study we define ILC as live CD45^+^ Lin^-^CD127^+^ (Figure 1A) and use KLRG1 as a key marker for intestinal lamina propria (LP) ILC2. NKp46 (encoded by the *NCR1* gene) has been well characterized to be expressed by ILC1 and NKp46^+^ ILC3 (reviewed by Vivier *et al.*, 2018). Surprisingly, we found a small population of NKp46-positive colonic LP (cLP) ILC2 in RORγt reporter mice (RORγt-eGFP^+/-^), and 64% of these cells did not express RORγt (Figure 1B). NKp46^+^ ILC2 could also be found in C57BL/6 mice, and approximately one third of these cells expressed ST2 (Figure 1C). Interestingly, NKp46^+^ cLP ILC2 were not present in mice deficient for T-bet suggesting a role for T-bet in their generation or maintenance (Figure 1C-D). IFNγ and IL-27 are known to induce T-bet expression. However, the genetic absence of either *Ifng* or *Il27ra (wsx1*) had no impact on the presence of NKp46^+^ cLP ILC2 (Fig 1E-F) suggesting that their roles are not critical for the generation or maintenance of these cells. As observed on the WT background, *Tbx21^-/-^Rag2^-/-^* ulcerative colitis (TRnUC) resulted in the near-complete loss of NKp46^+^ cLP ILC2 (Figure 1G-H).

**Figure 1.**
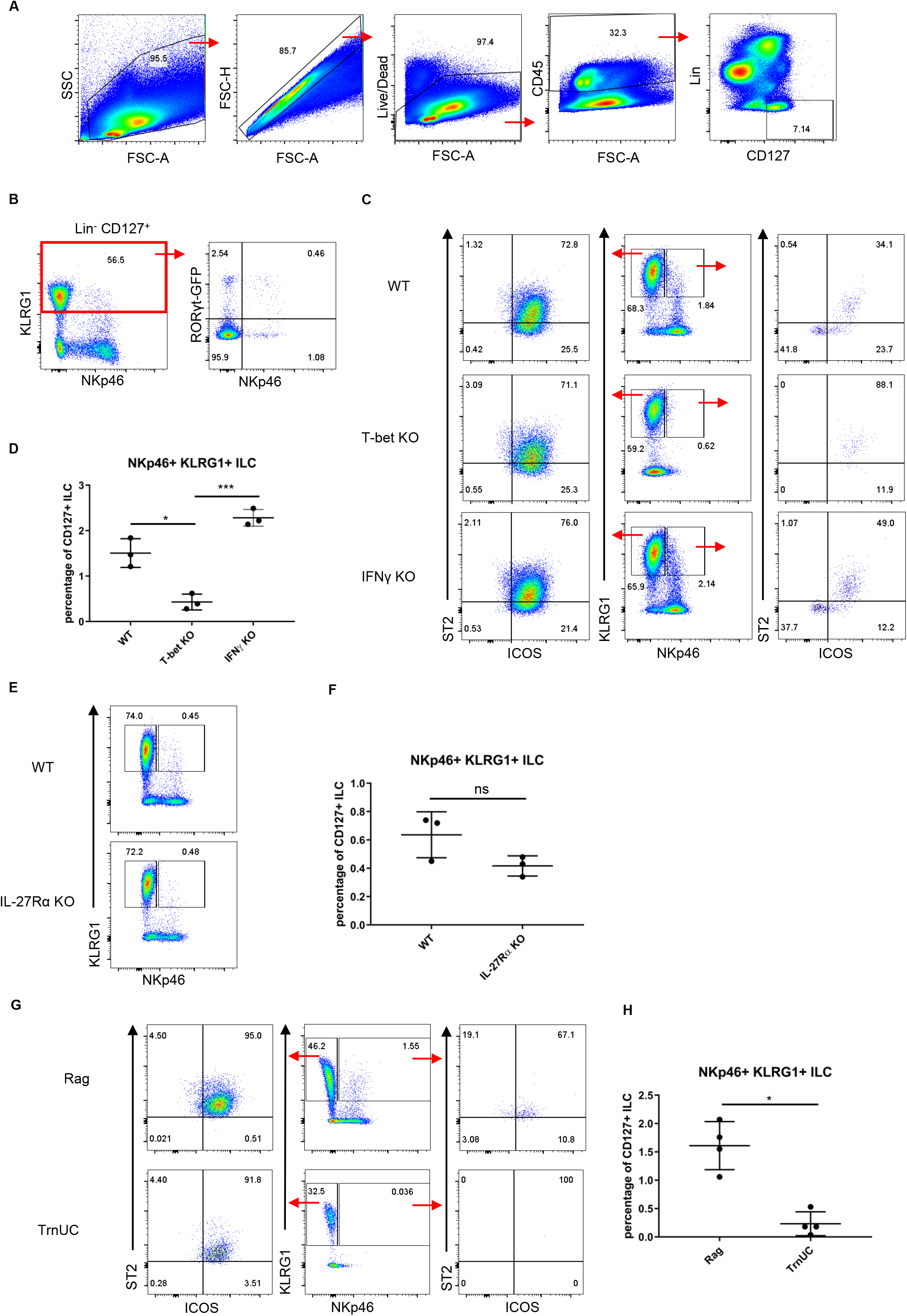
ILC2-like cells expressing NKp46 and T-bet exists in the intestinal lamina propria. ILC were isolated from the intestinal lamina propria for FACS analysis. (a) ILC were gated as live CD45^+^ Lin^-^ CD127^+^ leukocytes. (b) Expression of RORγt-GFP and surface NKp46 in live CD45^+^ Lin^-^ CD127^+^ ICOS^+^ KLRG1^+^ cLP ILC. Surface expression of NKp46, ICOS and ST2 in live CD45^+^ Lin^-^ CD127^+^ ICOS^+^ KLRG1^+^ cLP ILC in (c) C57BL/6, T-bet KO and IFNγ KO mice (e) C57BL/6 and IL-27Rα KO mice and (g) Rag and TrnUC mice, and (d, f, h) statistical analysis of NKp46^+^ KLRG1^+^ cLP ILC presence in these mice. Data shown are representative of a minimum of 3 biological replicates.

### A subset of intestinal ILC2 expresses T-bet

Further investigations into the existence of a T-bet-dependent population of cLP ILC2 showed that a small population of cLP ILC2 were found to actively express T-bet using a T-bet reporter mouse (Figure 2A, Figure 2D). Interestingly, many but not all T-bet^+^ ILC2 expressed NKp46 and ST2 expression in these cells was virtually absent. T-bet expression in cLP ILC2 may be temporal and not all ILC2 may be able to induce its expression. In order to detect cells with a history of T-bet expression, we generated a mouse model that expresses Cre-recombinase under the expression of the T-bet endogenous promoter by inserting an IRES-Cre cassette downstream of the *Tbx21* stop codon (Supplementary Figure 1A). This T-bet^Cre^ mouse was then bred to the Rosa26-lox-stop-lox YFP mouse (Rosa26^YFP/+^) (Srinivas *et al.*, 2001) to generate the T-*bet^Cre/+^xRosa26^YFP/+^* fate-mapper mouse (T-bet^FM^). As expected, cLP ILC1 defined as NKp46^+^NK1.1^+^T-bet^+^ were found to be T-bet fate mapper positive (T-bet^FM+^), while CCR6^+^ cLP ILC did not show expression of the fate mapper (Supplementary Figure 1B-C), confirming the functionality of the model – as these cells have not previously been shown to express T-bet. When examining ILC2 in these mice, we found that a subpopulation of T-bet^FM+^ ILC2 could be found within the lamina propria in the colon, small intestine and cecum with co-expression of GATA3 (Figure 2B, Figure 2D). Interestingly, in these ILC2, T-bet-YFP expression appeared to be predominately co-expressed with low expression of GATA3, but was also found in GATA3^hi^ expressing cells. In the T-bet reporter and FM model, approximately 5 % of intestinal lamina propria ILC2 demonstrated T-bet expression, suggesting that the same population of cells was being detected by both models. Although T-bet is known to be expressed by two known subsets of CD127^+^ ILC (ILC1 and NKp46^+^ ILC3), it was previously less appreciated that ILC2 cells may express this transcription factor as well at steady state.

**Figure 2.**
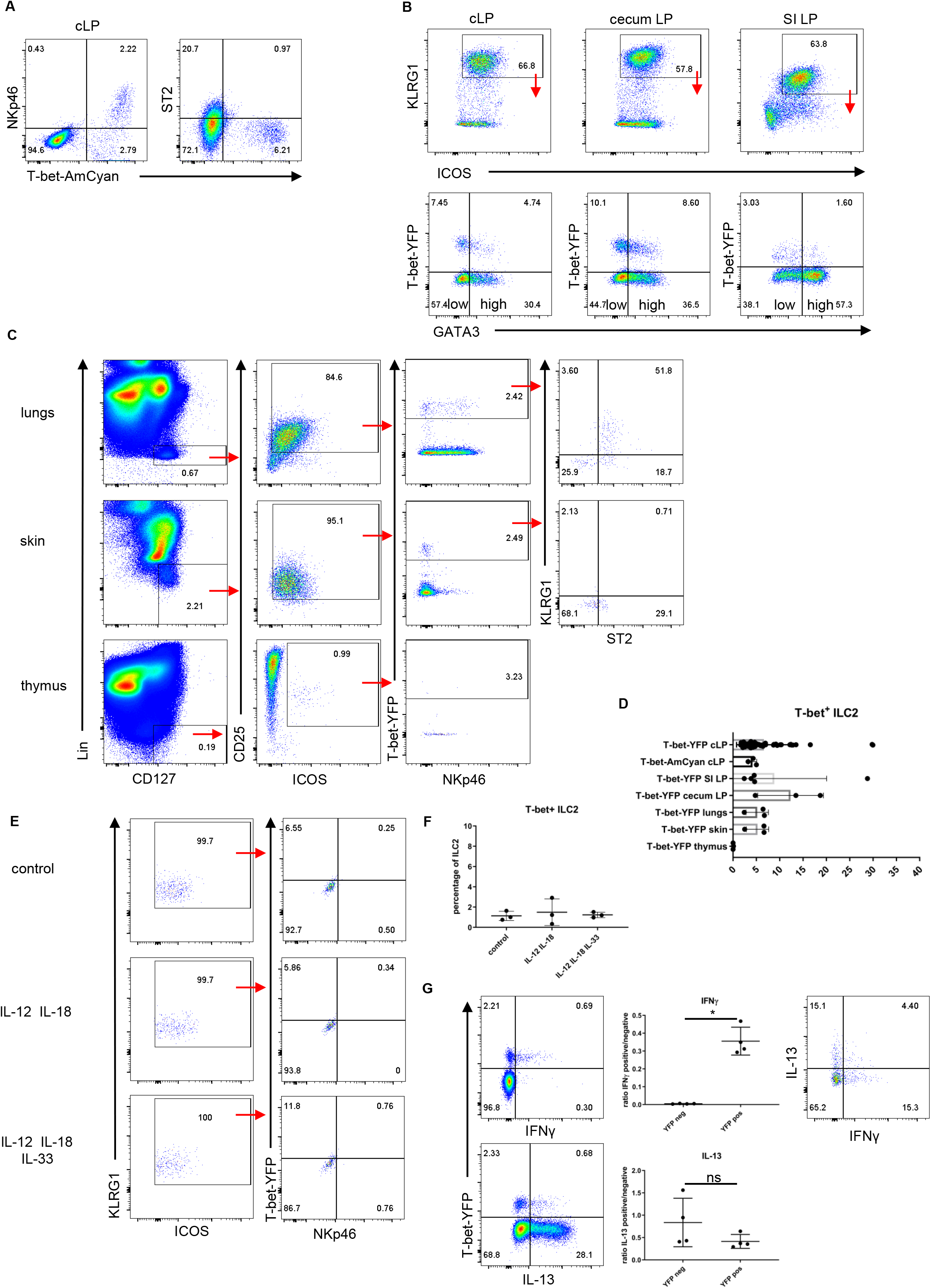
A subset of ILC2 expresses T-bet. ILC were isolated from various tissues for FACS analysis. (a) Surface expression of T-bet-AmCyan and ST2 in live CD45^+^ Lin^-^ CD127^+^ ICOS^+^ KLRG1^+^ cLP ILC in T-bet reporter mice. (b) T-bet fate mapper and GATA3 expression in live CD45^+^ Lin^-^ CD127^+^ ICOS^+^ KLRG1^+^ cLP, cecum LP and SI LP ILC2 in T-bet^FM^ mice. (c) T-bet fate mapper, ST2 and KLRG1 expression in live CD45^+^ Lin^-^ CD127^+^ ICOS^+^ CD25^+^ pulmonary, cutaneous and thymic ILC2 in T-bet^FM^ mice. (d) Summary plot for Figure 2a,b and c to show percentage of T-bet^+^ ILC2. (e,f) Rag2^-/-^ cLP ILC2 were sorted as live CD45^+^ Lin^-^ CD127^+^ ICOS^+^ KLRG1^+^ and cultured *in vitro* for 3 days in the presence of IL-7 and IL-2. Cytokines indicated were also added to the medium, and T-bet and NKp46 expression in live CD45^+^ Lin^-^ CD90.2^+^ ICOS^+^ KLRG1^+^ ILC was examined by FACS analysis after 3 days in culture and (f) percentage of T-bet expressing ILC2 are shown. (g) IL-13 and IFNγ expression and co-expression in live CD45^+^ Lin^-^ CD127^+^ KLRG1^+^ICOS^+^ T-bet^FM+^ ILC was examined by FACS analysis. Data shown are representative of a minimum of 3 biological replicates.

T-bet^FM+^ ILC2 defined as Lin^-^ CD127^+^ ICOS^+^ CD25^+^ ILC could further be identified in lungs and cutaneous tissue from ears, but not in the thymus (Figure 2C-D). In contrast to cutaneous T-bet^FM+^ ILC2, pulmonary T-bet^FM+^ ILC2 predominately showed expression of ST2 and KLRG1. Induced T-bet expression in ILC2 has been reported by a number of groups. The phenomenon is best described for human blood ILC2, but Silver *et al.* (2016) have confirmed that murine pulmonary ILC2 can also express T-bet and IFN-γ upon stimulation with IL-12 and IL-18 *in vitro*. However, using similar *in vitro* culture conditions, we were not able to induce T-bet and NKp46 expression in FACS purified cLP ILC2 from Rag2-deficient mice upon stimulation with IL-12 and IL-18 and stimulation with IL-12, IL-18 and IL-33 had no effect either (Figure 2E-F). Hence, it may be possible that plasticity in ILC2 and generation of T-bet^+^ ILC2 may differ mechanistically depending on their tissue location.

In order to characterise the potential effector functions of T-bet^FM+^ cLP ILC2, we analysed their ability to express the T-bet target cytokine IFNγ alongside IL-13, conventionally expressed by ILC2. Remarkably, these cells did express IFNγ in contrast to conventional ILC2 while maintaining ability to express IL-13, while T-bet^FM-^ ILC2 expressed only IL-13. However, T-bet^FM+^ ILC2 demonstrated only limited co-expression of these cytokines (Figure 2G).

### T-bet^FM+^ cLP ILC2 show a distinct genetic profile

To further characterise T-bet^FM+^ and T-bet^FM-^ ILC2, differentially expressed transcripts were analysed using FACS purified cLP cells. T-bet^FM+^ ILC2 shared a gene expression profile with T-bet^FM-^ ILC2 in relation to ILC2-signature genes, while striking differences were also identified (Figure 3A). The expression of transcripts for the ILC2-related cytokines IL-13, IL-5 and IL-4, was equivalent in T-bet^FM+^ and T-bet^FM-^ ILC2, as was expression of amphiregulin (*Areg).* In addition, both T-bet^FM+^ and T-bet^FM-^ ILC2 expressed transcripts for the IL-33 and IL-25 receptors subunits (*Il1rl1* (ST2) and *Il17rb* respectively) and the signature ILC2 surface markers KLGR1 and ICOS. Similarly, the transcription factor RORα was homogeneously expressed in both populations. These results confirm that this T-bet^FM+^ population expresses the core transcriptional profile associated with ILC2. Of note, neither T-bet^FM+^ nor T-bet^FM-^ ILC2 expressed genes related to an NK cell-genetic profile (*Eomes, Gzmb, Prf1* or *Tnfsf10*).

**Figure 3.**
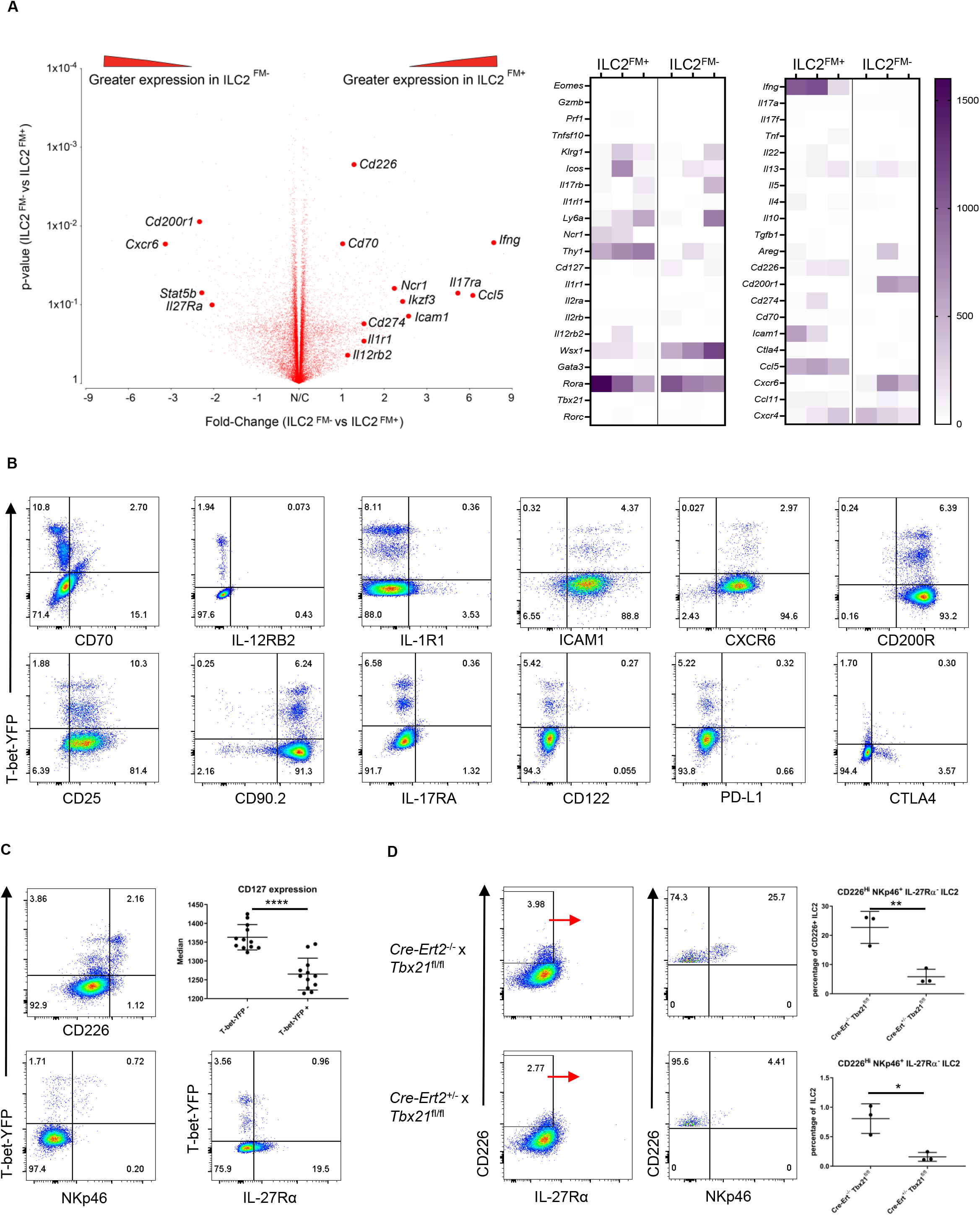
ILC1-ILC2 show a distinct genetic profile and dependent on T-bet. T-bet^FM+^ and T-bet^FM-^ cLP ILC2 were FACS sorted to perform a genetic profiling. (a) Volcano plot comparing the fold change in gene expression in T-bet^FM+^ and T-bet^FM-^ cLP ILC2 isolated from T-bet^FM^ mice. A heat map shows the expression of selected genes in T-bet^FM+^ and T-bet^FM-^ cLP ILC2. cLP ILC2 from T-bet^FM^ mice were isolated for FACS analysis. (b) IL-12RB, PD-L1 (CD274), ICAM-1, CD122, CD90.2, CXCR6, IL-17RA, CD70, CTLA4, IL-1R1 and CD200R expression in live CD45^+^ Lin^-^ CD127^+^ ICOS^+^ KLRG1^+^ T-bet^FM^ cLP ILC. (c) NKp46, CD226, IL-27Rα and CD127 expression in live CD45^+^ Lin^-^ CD127^+^ ICOS^+^ KLRG1^+^ T-bet^FM^ cLP ILC. (d) CD226, NKp46 and IL-27Rα expression on live CD45^+^ Lin^-^ CD127^+^ ICOS^+^ KLRG1^+^ cLP ILC from tamoxifen pre-treated WT and Het Cre-Ert2-T-bet^fl/fl^ mice were analysed by FACS. Data shown are representative of a minimum of 3 biological replicates.

*Tbx21* transcripts were not detected in either T-bet^FM-^ ILC2 or T-bet^FM+^ ILC2, but T-bet^FM+^ ILC2 expressed *Ncr1* (encoding NKp46) and the T-bet target gene *Ifng.* Similarly, the expression of other T-bet related genes, such as *Cd226*, *Il1r1* and *Il12rb2* was up-regulated in T-bet^FM+^ ILC2. Other genes were also differentially expressed between T-bet^FM+^ and T-bet^FM-^ ILC2, including *Cxcr6, Cd70, Wsx1, Cd220r, Il17ra, Thy1, Icam* or *Ccl5*, highlighting the differences between these two ILC2 subsets. Therefore, T-bet^FM+^ ILC2 display a unique transcriptional profile, supporting the hypothesis that T-bet^FM+^ constitute an ILC subset distinct from ILC2. Hence from hereon, we refer to these cells as ILC1-ILC2.

To further characterize ILC1-ILC2, the expression of candidate surface markers identified by the gene array analysis was assessed on cLP ILC1-ILC2 and ILC2 (Figure 3B-C). Corroborating the gene array data, the ILC1-ILC2 population had greater expression of NKp46 and CD226 (DNAM-1) than conventional cLP ILC2 (Figure 3C). However, not all ILC1-ILC2 expressed NKp46 or high levels of CD226 indicating that these cells are heterogeneous. Both CD226 and NKp46 are known to be expressed by ILC1 and NK cells, and their expression has been correlated with T-bet expression (reviewed by Chiossone *et al.*, 2018). CD226 binds to CD112 and CD155 on transformed cells resulting in NK cell-mediated lysis and IFN-γ production from NK cells (Tahora-Hanaoka *et al.*, 2004). Likewise, the ligation of NKp46 expressed by NK cell to its ligand results in cytotoxicity and cytokine production (Glasnet *et al.*, 2018, Lakshmikanth *et al.*, 2009, Halfteck *et al.*, 2009). In contrast to CD226 and NKp46, some cLP ILC2 showed surface expression of the IL-27R subunit IL-27Rα (Wsx-1), while cLP ILC1-ILC2 did not express this receptor as predicted from the RNA array. IL-27 signalling has been reported to inhibit IL-13 and IL-5 expression by lung ILC2 and Th2 cells (Moro *et al.*, 2016, Stumhofer *et al.*, 2007). In addition, IL-27 has been suggested to induce T-bet in T cells (Hibbert *et al.*, 2003). The absence of surface IL-27R on cLP ILC1-ILC2 indicates that these cells may be regulated differently from conventional cLP ILC2, at least at steady state. Furthermore, ILC1-ILC2 had a lower expression of CD127 in comparison to conventional ILC2 (Figure 3C), and this is in line with previous publications indicating that T-bet restricts CD127 expression in cLP ILC (Garrido Mesa *et al.*, 2019, Powell *et al.*, 2012). In contrast, surface expression of IL-12RB2, PD-L1 (CD274), ICAM-1, CD122, CD90.2, CXCR6, IL-17RA, CD70, CD200R, CD25, IL-1R1 and CTLA4 was similar between ILC1-ILC2 and ILC2 (Figure 3B).

*Tbx2^fl/fl^* Cre-Ert2 ^+/-^ (Het) and *Tbx2 Cre-Ert2^-/-^* (WT) mice were employed to test the post-developmental requirement of T-bet for CD226^Hi^ NKp46^+^ ILC1-ILC2 maintenance. Tamoxifen was injected into these mice each day for a period of 5 days. 3 weeks after the first injection we observed a significant loss of cLP ILC1-ILC2 expressing NKp46 and high levels of CD226 (Figure 3D). This highlights the critical role of T-bet for the maintenance of these cells.

### Functional analysis of cLP ILC1-ILC2

We have previously shown that T-bet deficiency on a C57BL/6 background drives a more potent immune response to *T. spiralis* (Alcaide *et al.*, 2007, Garrido Mesa *et al.*, 2019), a phenomenon attributed to enhanced type 2 immunity. Hence we explored whether T-bet-expressing ILC, including ILC1-ILC2, may play a functional role in this model in the absence of T and B cells. However, the worm burden of *T. spiralis* was not altered in TRnUC mice when compared with T-bet sufficient animals, indicating that lack of T-bet expression in the innate immune system plays no functional role in this model (Supplementary Figure 2A). In addition, ILC1-ILC2 are also unlikely to play any significant role in the immune response to the intestinal helminths *N. brasiliensis* or *H. polygyrus*. For these experiments, the role of T-bet was tested in mice with a germline depletion of T-bet and also in the model of induced depletion of T-bet using Cre-Ert2^-/+^ mice. In none of these models was it possible to record an altered anti-parasite host response in the absence of T-bet. Fecal egg counts or worm burden in the small intestine was not altered upon infection and there were no differences between the mouse models for goblet cell counts as determined by Periodic acid-Schiff (PAS) stain, throughout infection time course (Supplementary Figure 2B-I).

In order to identify a functional role for ILC1-ILC2 we utilised a dextran sulphate sodium (DSS) model of induced colitis in T-bet^FM^ mice. These mice developed acute colitis 2 days after the withdrawal of DSS (Figure 4A). Although ILC1-ILC2 were potent producers of IFNγ and IL-13 in the fresh water control as outlined above, these cytokine-producing cells were significantly diminished in mice with DSS-induced colitis (Figure 4B). This finding indicates that ILC1-ILC2 may play a role in the intestinal immune response in colitis that is distinct from that of ILC1 or ILC2.

**Figure 4:**
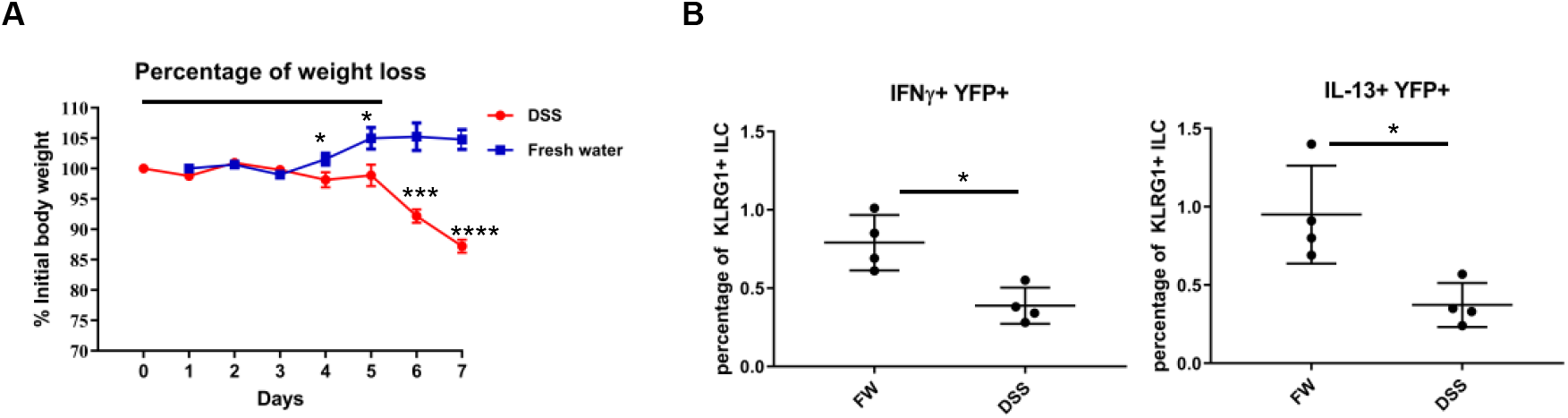
ILC-ILC2 produce less IFNγ and IL-13 during DSS-induced colitis. T-bet^FM^ mice were treated with 3% DSS in the drinking water for 5 days, and cLP ILC were harvested for FACS analysis after another 2 days. Control mice received fresh water (FW) at the same time. (a) Analysis of body weight loss in mice receiving DSS or fresh water is demonstrated. (b) Percentage of IL-13 and IFNγ expressing live CD45^+^ Lin^-^ CD127^+^ KLRG1^+^ ICOS^+^ T-bet^FM^ ILC was examined by FACS analysis. Data shown are plotted from 4 biological replicates.

### ILC1-ILC2 are unlikely to derive from ILC2 precursors

The origin of intestinal ILC1-ILC2 could be a separate developmental pathway or the result of active plasticity of ILC. To explore the possibility that ILC1-ILC2 are a separate novel ILC subset, we first analysed these cells in the lamina propria of colons and small intestines of mice at the age of weaning (3 weeks post-partum). ILC1-ILC2 were indeed present in both of these tissues (Figure 5A, Supplementary Figure 3A). These cells did not express α4β7 indicating they did not migrate into the tissue recently. Furthermore, we could only detect very few T-bet^FM+^ ILC2p and T-bet^FM+^ CLP populations in the bone marrow of some, but not all, adult mice (Figure 5B, Supplementary Figure 3B). ILC2p were defined as Lin^-^ CD127^+^ α4β7^hi^ Flt3^-^ CD25^+^ while CLP were gated as Lin^-^ CD127^+^ α4β7^-^ Flt3^+^ CD25^-^ bone marrow cells. Hence, we sought to determine whether we could generate T-bet^FM+^ ILC2 from T-bet^FM-^ CLP or ILC2p in a 6 day culture on OP9-DL1 stromal cells in the presence of rIL-7,rSCF and rIL-33 using an established method to develop ILC2. We were able to generate ILC2 from either source, but it was not possible to detect T-bet^FM+^ ILC2 in these cultures (Figure 5C, Supplementary Figure 3C). This may indicate that ILC1-ILC2 develop in the periphery or the tissue after extravasation, or from an undefined precursor or pathway. As we were not able to detect T-bet^FM+^ ILC2p consistently perhaps due to the rarity of these cells, we sorted this population from the pooled bone marrow of multiple animals and cultured them on OP9-DL1 cells. Strikingly, these T-bet^FM+^ ILC2p differentiated into T-bet^FM+^ ILC2 (Supplementary Figure 3D). Hence, it is plausible that T-bet^FM+^ ILC2p are precursors of ILC1-ILC2. Interestingly, when we analysed ILC1-ILC2 in one-week old litters, we could not detect ILC1-ILC2 in the colonic lamina propria (Figure 5D). α4β7^-^ ILC2 were present in both colonic and small intestinal lamina propria, however, ILC1-ILC2 could only be found in a subpopulation of KLRG1^+^ ICOS^low^ ILC2 present in SI LP, but not cLP (Figure 5D-E). These SI LP ILC1-ILC2 expressed α4β7 indicating that these cells may have emigrated recently from the bone marrow. Therefore, we interrogated an ILC2 specific single cell RNA-seq data set (Ricardo-Gonzalez *et al.*, 2018) and were able to detect T-bet^+^ ILC2 in a specific subpopulation of bone marrow cells (Figure 5F, Supplementary Figure 3E-F).

**Figure 5.**
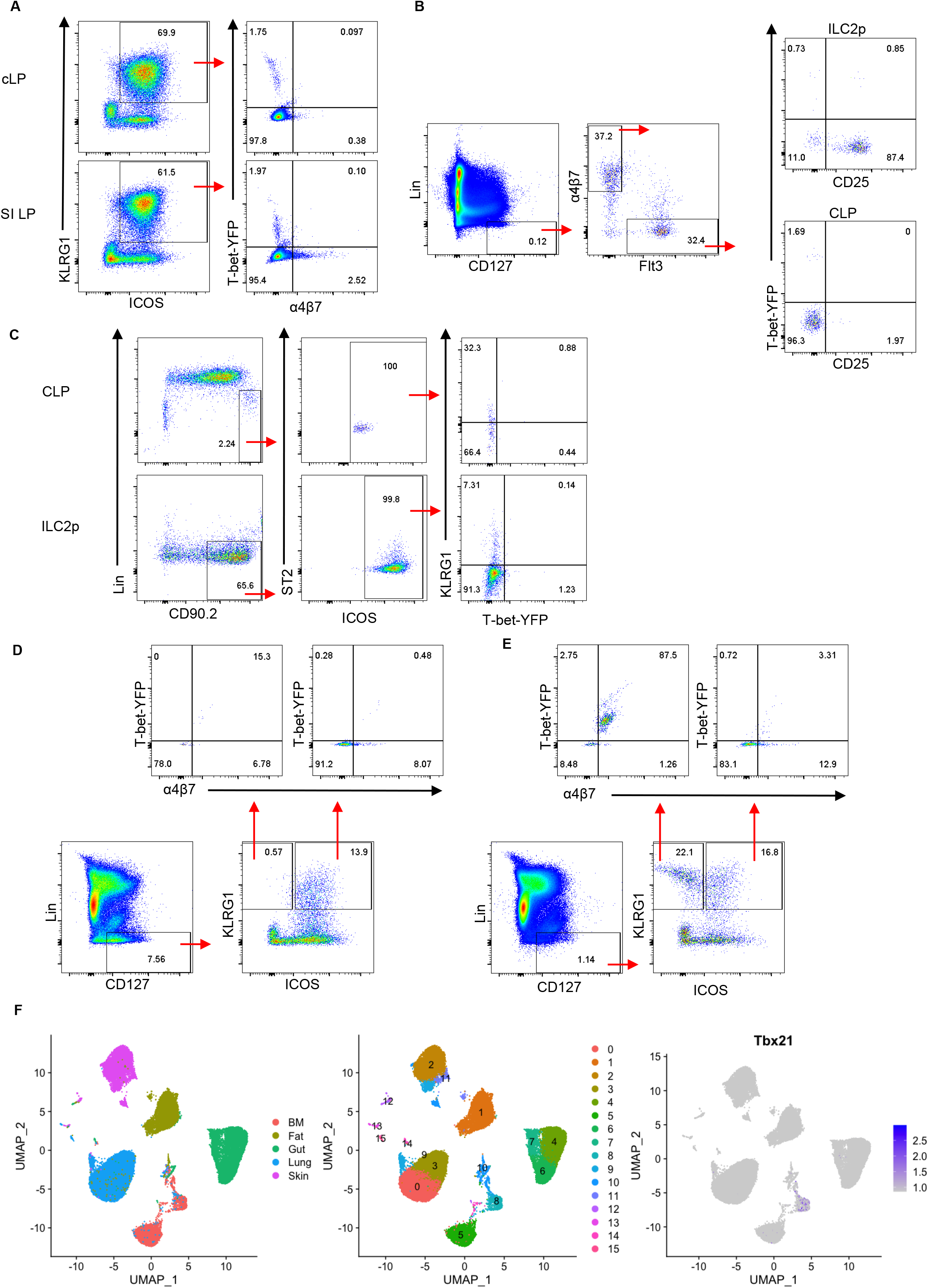
ILC1-ILC2 do not derive from a T-bet^FM-^ ILC2p. ILC were examined by FACS analysis. (a) T-bet fate mapper expression and α4β7 expression in live CD45^+^ Lin^-^ CD127^+^ ICOS^+^ KLRG1^+^ T-bet^FM^ cLP and SI LP ILC in 3 weeks old T-bet^FM^ mice. (b) T-bet fate mapper expression in live CD45^+^ Lin^-^ CD127^+^ α4β7^+^ CD25^+^ bone marrow ILC2p and live CD45^+^ Lin^-^ CD127^+^ Flt3^+^ bone marrow CLP in T-bet FM mice. (c) Generation of live CD45^+^Lin^-^ CD90.2^+^ ICOS^+^ KLRG1^+^ ILC from T-bet^FM-^ CLP or T-bet^FM-^ ILC2p in an OP9-DL1 co-culture system. T-bet fate mapper expression in live CD45^+^ Lin^-^ CD127^+^ ICOS^+^ KLRG1^+^ is presented. (d) cLP and (e) SI LP ILC in one-week old T-bet^FM^ mice. Data shown are representative of a minimum of 3 biological replicates (a-c) or shows one sample pooled from tissues of 7 mice (d-e). (f) A published ILC2-specific single cell RNA-seq dataset (Ricardo-Gonzalez *et al.*, 2018) was used to identify T-bet-expressing ILC2 in multiple organs. A cluster map showing the prevalence of T-bet^+^ ILC2 in various organs is shown.

This study is the first report of a novel murine subset of intestinal LP ILC1-ILC2 *in vivo.* Our data indicate a functional, genetic and protein expression profile of ILC1-ILC2 cells that is distinct from conventional intestinal LP ILC2, ILC1, ILC3 and NK cells.

## METHODS

### Animals

*Rosa26^YFP/+^* (Jackson labs) mice were sourced commercially and bred with T-bet^cre/+^ mice to generate the T-bet^Cre/+^x*Rosa26*^YFP/+^ (T-bet^FM^) mice. All mice were housed in specific pathogen–free facilities at King’s College London Biological Services Unit, London School of Hygiene and Tropical Medicine Biological Service Facility, University of Manchester Biological Services Facility or at Charles River Laboratories. T-bet-AmCyan and RORγt reporter mice were a gift from Dr Jinfang Zhu (Yu *et al.*, 2015) and Dr Gérard Eberl (Lochner *et al.*, 2011), respectively. C57BL/6 WT (Charles River), T-bet KO, IFN-γ KO, IL-27Rα KO (Charles River) and Ert2 (Jackson labs) mice were sourced commercially (all C57BLl/6 background). *T-be^fl/fl^* mice were previously generated by our group (Gökmen *et al.*, 2013) and crossed with Ert2 mice to generate *Cre-Ert2^+/-^* (Het) and *Cre-Ert2^-/-^* (WT) mice expressing *T-bet^fl/fl^*. A colony of colitis-free Rag2^-/-^xTbx21^-/-^ (TRnUC) mice was generated from the TRUC colony that was descendant from the originally described TRUC mice (Powell *et al.*, 2012).

#### Generation of T-bet^cre/+^ mouse

To allow the expression of the Cre-recombinase under the expression of the T-bet endogenous promoter, a T-bet knock-in mouse was generated (GenOway, France). For this purpose, an IRES-Cre cassette was introduced downstream of the Stop codon of the T-bet gene, in the 3’UTR (Supplementary Figure S1A). The genomic region of interest containing the murine Tbx21 locus was isolated by PCR from 129Sv genetic background. PCR fragments were subcloned into the pCR4-TOPO vector (Invitrogen). The genomic clones (containing intron 1 to exon 6) were used to construct the targeting vector. Briefly, a 5.6-kb fragment comprising Tbx21 exon 2 and 6 and a 1.6-kb fragment located downstream of the Tbx21 exon 6 STOP codon were used to flank an IRES-Cre cassette (FRT site-PGK promoter-Neo cDNA-FRT site).

#### Screening of T-bet–targeted embryonic stem cell clones

The FseI-linearized targeting vector was transfected into C57BL/6 ES cells. Positive selection was started 48 hours after electroporation, by addition of 200 μg/ml G418 (150 μg/ml active component; Life Technologies). Then, 275 resistant clones were isolated, amplified, and screened by PCR and further confirmed by Southern blot.

#### Generation of chimeric mice and breeding scheme

One floxed mutated *Tbx21* ES cell clone was microinjected into albino C57BL/6 strain (C57BL/6J-Tyrc-2J/J) blastocysts, and gave rise to male chimeras with a significant ES cell contribution (as determined by the percentage of light and dark patches on their coat). After mating with C57BL/6 CMV-Flp–expressing female mice to remove the FRT-flanked Neo cassette, offspring were genotyped by PCR and Southern blot to ensure removal of the Neo cassette. PCR and Southern blot screening conditions are available on request. The mosaic excised F1 mouse was them mated with C57BL/6 WT mice to obtain a pure line of Cre-expressing T-bet knock-in mice: T-bet^cre/+^.

#### Tamoxifen induction of Tbx21 depletion

A tamoxifen stock solution was prepared by suspending 1g of tamoxifen (free base: MP biomedicals, LLC) into 18ml 100% ethanol and heating to 37°C briefly to dissolve the tamoxifen. Prior to the injections this solution was diluted at 1:4.56 in sunflower oil and heated briefly to 37°C. *In vivo* depletion of T-bet was induced by consecutive intraperitoneal injections of tamoxifen on five consecutive days with one injection per day (1mg per injection). Three weeks after the first injection of tamoxifen tissues were harvested or the mice were infected or treated with DSS as outlined below.

### Isolation of cells

cLP, SI LP and cecum LP leukocytes were isolated using a published method (Grohnke *et al.*, 2017). Peyer’s patches were removed from SI prior to cell isolation. Briefly, the epithelium was removed by incubation in HBSS lacking Mg^2+^ or Ca^2+^ (Invitrogen) supplemented with EDTA and HEPES. The tissue was further digested in 2% of foetal calf serum (FCS Gold, PAA Laboratories) supplemented in 0.5mg/ml collagenase D, 10μg/ml DNase I and 1.5mg/ml dispase II (all Roche). The LP lymphocyte-enriched population was harvested from a 40%-80% Percoll (GE Healthcare) gradient. Lung leukocytes were cut in small pieces and cells were further isolated by an incubation in 300IU/ ml collagenase II (Gibco) and 3μg/ml DNase I (Roche) for one hour at 37 degree C. Ear sheets were incubated in 5 mg/ ml collagenase IV (Sigma) for one hour at 37 degree C. Cutaneous leukocytes were further purified by Percoll gradient application. Thymic cells were isolated as described before (Schroeder *et al.*, 2013).

### Flow Cytometry

Flow cytometry was performed using a standard method. For ILC stains a lineage cocktail was used consisting of antibodies specific for CD3, B220, CD19, CD11b, Ter-119, Gr-1, CD5 and FcεRI. In case of thymic tissue antibodies specific for CD4 and CD8 were added to the lineage mix, and the lineage mix for skin tissue contained anti-CD2 antibodies. For a complete list of the antibodies used see Table 1. Samples were acquired using an LSRFortessa™ cell analyser (Becton Dickinson, USA), and data were analysed using FlowJo software (Tree Star, USA). A FoxP3 staining kit (Invitrogen) was used for staining CTLA4, cytokines and transcription factors. In case of cytokine analysis, cells were pre-stimulated with 100 ng/ml PMA and 2μM ionomycin in the presence of 6μM monensin for 3 hours prior to FACS analysis at 37°C 5% CO2.

**Table 1:** Antibody clones and distributors

| **Antibody** | **Clone** | **Company** |
| --- | --- | --- |
| CD3 | 17A2 | eBioscience |
| CD5 | 53-7.3 | eBioscience |
| CD19 | 1D3 | eBioscience |
| B220 | RA3-6B2 | eBioscience |
| CD11b | M1/70 | eBioscience |
| Gr-1 | RB6-8C5 | eBioscience |
| Ter119 | TER-119 | eBioscience |
| FcεRI | MAR | eBioscience |
| CD127 | A7R34 | eBioscience |
| NKp46 | 29A1.4 | eBioscience |
| ST2 | RMST2-33 | eBioscience |
| ICOS | C398.4 | eBioscience |
| KLRG1 | 2F1 | eBioscience |
| CCR6 | 29-2L17 | eBioscience |
| GATA3 | L50-823 | BD |
| NK1.1 | PK136 | Biolegend |
| IL-13 | eBio13A | eBioscience |
| IFNγ | XMG1.2 | eBioscience |
| T-bet | 4B10 | Invitrogen |
| CD45 | 30-F11 | Invitrogen |
| $\alpha 4\beta 7$ | DATK32 | eBioscience |
| CD70 | FR70 | Biolegend |
| IL-12RB2 | 305719 | R&D |
| IL-1R1 | JAMA-147 | eBioscience |
| ICAM-1 | YN1/1.7.4 | eBioscience |
| Flt3 | A2F10 | eBioscience |
| CXCR6 | DANID2 | Invitrogen |
| CD200R | OX110 | eBioscience |
| CD25 | PC61.5 | eBioscience |
| CD90.2 | 30-H12 | BD |
| IL-17RA | PAJ-17R | eBioscience |
| CD122 | TM-β1 | eBioscience |
| PD-L1 | MIH5 | eBioscience |
| CTLA4 | UC10-4B9 | Invitrogen |
| IL-17Ra | 2918 | BD |
| CD226 | 10 E5 | eBioscience |
| CD2 | RM2-5 | eBioscience |
| CD4 | RM4-5 | eBioscience |
| CD8 | 53-6.7 | eBioscience |

### Cell Sorting and *in vitro* culture

Single-cell suspensions from the intestine or bone marrow were stained with fluorescently labelled antibodies and sorted (purity > 98%) using a BD FACSAria III cell sorter (BD Biosciences). For ILC, antibodies against CD45, lineage markers and IL-7Rα and DAPI stain were used to separate live CD45^+^Lin^-^IL-7Rα^+^ cells. cLP Rag2^-/-^ ILC2 (ICOS^+^ KLRG1^+^) were cultured in DMEM supplemented with 10% FCS, 1xGlutaMax (Gibco), 50 U/ml penicillin, 50 μg/ml streptomycin, 10 mM HEPES, 1x non-essential amino acids (Gibco), 1 mM sodium pyruvate and 50 μM β-mercaptoethanol (Gibco). The medium was further supplemented with rmIL-7 and rhIL-2 (both at 10 μg/ml) and further cytokines as indicated (all cytokines were used at a final concentration of 10 μg/ml). Cells were harvested and analysed by flow cytometry after 3 days in culture at 37°C 5% CO2.

### ILC generation in OP9-DL1 system

CLP or ILC2p were seeded on OP9-DL1 to generate ILC2 using an established method (Seehus *et al.*, 2016). Briefly, 7,500 cells were co-cultured with mitomycin pre-treated OP9-DL1 in presence of rmIL-7, rmSCF and rmIL-33 (all 20 ng/ml) for 6 days prior to FACS analysis.

### Gene expression array

RNA was extracted using Trizol reagent (Invitrogen, UK), according to the manufacturer’s protocol and contaminating DNA was removed with the RNase-Free DNase Set (Qiagen, UK).

cDNA was synthetized using Ovation One-Direct System and labelled using Encore BiotinIL module (Nugen, USA) following the manufacture’s protocol. RNA and cDNA quantity and quality were assessed using the Agilent RNA 6000 Nano Kit and Agilent RNA 6000 Pico Kit according to the manufacture’s protocol (Agilent Technologies, USA). Labelled cDNA were hybridised on a MouseWG-6 v2.0 Expression BeadChip (Illumina, USA) and gene expression analysis was performed using Partek software (Partek Incorporated, USA). All raw and processed gene expression array data are available at Gene Expression Omnibus (www.ncbi.nlm.nih.gov/geo/).

### Infection mouse models

Mice were infected with *T. spiralis, N. brasiliensis* or *H. polygyrus* using published methods (Turner *et al.*, 2013, Entwisle *et al.*, 2017, Garrido Mesa *et al.*, 2019). In some experiments T-bet depletion was induced prior to infection with *N. brasiliensis* or *H. polygyrus.*

### DSS-induced colitis

Colitis was induced by adding 3% DSS (36-50 KDa, MP Biomedicals, Ontario, USA) to the drinking water for 5 days. Non-colitic mice were administered sterile drinking water. Mice were sacrificed 7 days after the beginning of the experiment. Weight and clinical abnormalities were monitored on a daily basis.

### Single-cell RNA-seq analysis

Raw expression matrices were obtained from GEO (GSE117567, Ricardo-Gonzalez *et al.* 2018), then processed using Seurat 3.0 (Stuart et al 2019). Genes detected in less than 5 cells, and cells with less than 500 detected genes, were removed. Following normalization (using NormalizeData), samples were merged into a single object, and the top 2000 variable genes were used to calculate the PCA (PCs = 30), then UMAP (dims = 20) reductions. Shared nearest neighbour and clustering were carried out using FindNeighbours (reduction = PCA, dims = 20) and FindClusters (res = 0.5) respectively.

### Statistics

Results are expressed as mean ± SEM. Data were analysed using Student’s t-test or Mann-Whitney U test, as appropriate, using GraphPad Prism 5.0 (GraphPad Inc., USA). ns: non-significant; *p < 0.05; **p< 0.01; ***p<0.001; ****p<0.0001.

### Study approval

All animal experiments were performed in accredited facilities in accordance with the UK Animals (Scientific Procedures) Act 1986 (Home Office Licence Numbers PPL: 70/6792, 70/8127, 70/7869 and 60/4423).

## ACKNOWLEDGMENTS

We thank the members of the LORD laboratory for valuable discussions and critically commenting on the manuscript. In addition, we thank the BRC flow cytometry core team for technical help, and acknowledge financial support from the Department of Health via the NIHR comprehensive Biomedical Research Centre award to Guy’s and St. Thomas’ NHS Foundation Trust in partnership with King’s College London and King’s College Hospital NHS Foundation Trust. We also thank Professor Zúñiga-Pflücker (Sunnybrook Research Institute, University of Toronto) for contributing OP9-DL1 cells and Dr Jinfang Zhu (National Institute of Allergy and Infectious Diseases, USA) for providing T-bet-AmCyan mice.

## AUTHOR CONTRIBUTIONS

Study concept and design (JHS, NG, RGJ, PL, JKH, HH, RKG, JFN, GML), acquisition of data (JHS, NG, TZ, ALG, LC, ES, LBR, JWL, JFN), data analysis and interpretation (JHS, NG, PL, GB, LBR, JWL, JFN, GML), technical support (AI, CMH, RR), obtained funding (NG, JFN, JKH, GML), drafting of manuscript (JHS), study supervision (JFN, GML).

## DISCLOSURE

The authors have no conflict of interest to declare.

---

Supplementary Figure 1. Design and evaluation of T-bet fate mapper mouse

cLP ILC were isolated for FACS analysis. (a) See method section for further details on the design. (b) T-bet^FM^ and T-bet expression in live CD45^+^ Lin^-^ CD127^+^ NKp46^+^ NK1.1^+^ cLP ILC. (c) T-bet^FM^ expression in live CD45^+^ Lin^-^ CD127^+^ NKp46^+^ CCR6^-^ and NKp46^-^ CCR6^+^ cLP ILC. Data shown are representative of a minimum of 3 biological replicates.

Supplementary Figure 2. Intestinal parasite infection in in the absence of T-bet

(a) Rag and TRnUC mice were infected with *T. spiralis* for infection analysis. Worm counts in the muscles at day 9 post infection are shown. (b, d) C57BL/6 and T-bet KO or (c, e) tamoxifen pre-treated WT and Het Cre-Ert2-T-bet^fl/fl^ mice were infected with *N. brasiliensis* for infection analysis. (b, c) PAS-stained histology sections of infected SI tissue and goblet cell numbers per vili and are shown. (d, e) Feces egg counts of *N. brasiliensis* at day 5, 6, 7, 8 and 9 post infection and SI worm count of *N. brasiliensis* at day 7 and 9 post infection are presented. (f, h) C57BL/6 and T-bet KO or (g, i) tamoxifen pre-treated WT and Het Cre-Ert2-T-bet^fl/fl^ mice were infected with *H. polygyrus* for infection analysis. (f, g) PAS-stained histology sections of infected SI tissue and goblet cell numbers and are shown. (h, i) Feces egg counts of *H. polygyrus* at day 14 post infection and SI worm count of *H. polygyrus* at day 14 post infection are presented. Data shown are plotted from a minimum of 3 biological replicates.

Supplementary Figure 3. T-bet-positive ILC2 can be predominately detected in the bone marrow.

Summary plots for (a) Figure 5a, (b) Figure 5b and (c) Figure 5c to demonstrate T-bet expression. (d) FACS analysis of live CD45^+^ Lin^-^ CD90.2^+^ ICOS^+^ KLRG1^+^ ILC generated from T-bet^FM-^ and T-bet^FM+^ ILC2p in an OP9-DL1 co-culture system. T-bet fate mapper expression in live CD45^+^Lin^-^ CD127^+^ ICOS^+^ KLRG1^+^ is shown. Data shown are derived from BM cells pooled from nine mice. (e,f) A published ILC2-specific single cell RNA-seq dataset (Ricardo-Gonzalez *et al.*, 2018) was used to identify T-bet-expressing ILC2 in multiple organs. A distinct genetic profile of BM T-bet^+^ ILC2 is illustrated by comparing to the average expression levels of specific markers in ILC2 from all other organs.

## Notes

**Financial support:** This study was supported by grants awarded by the Wellcome Trust (GML, 091009) and the Medical Research Council (GL, TTM, MR/M003493/1; GML, JKH, MR/K002996/1). JFN was supported by a RCUK/UKRI Rutherford Fund fellowship (MR/R024812/1). Research was also supported by the National Institute for Health Research (NIHR) Biomedical Research Centre at Guy’s and St Thomas and King’s College London (GL). NG was funded by Fundación Ramón Areces (Spain) and British Heart Foundation (PG/12/36/29444). The views expressed are those of the author(s) and not necessarily those of the NHS, the NIHR, or the Department of Health.

### Competing Interest Statement

The authors have declared no competing interest.

