## Supplementary figures and images for "T-bet fate mapping identifies a novel ILC1-ILC2 subset *in vivo*"

### Supplementary files

#### Supplementary Figure 1

**A**

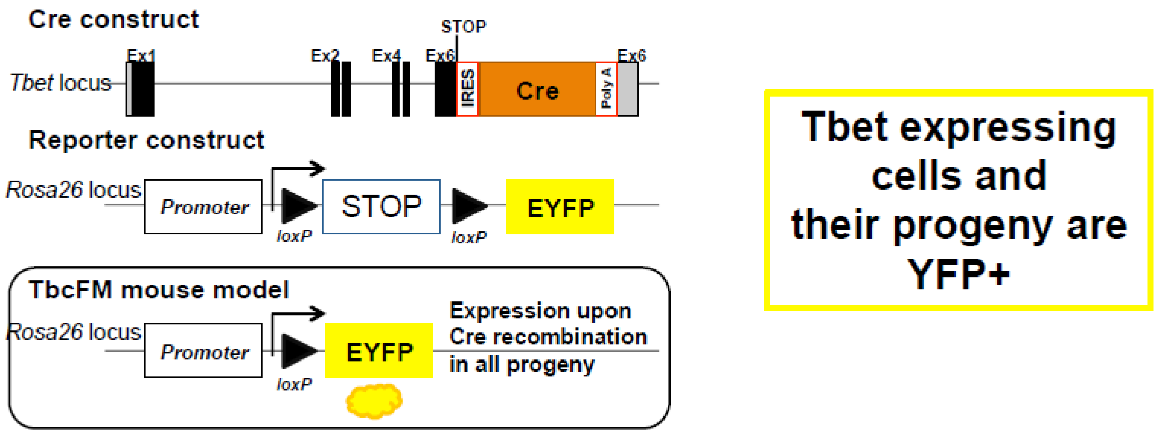


**B**

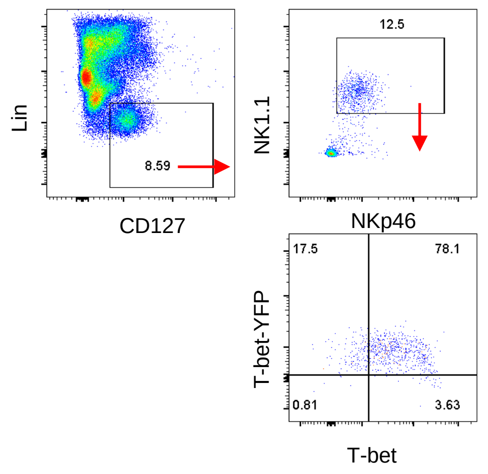


**C**

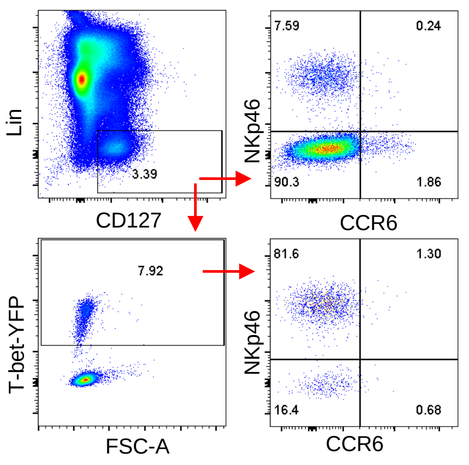

#### Supplementary Figure 2

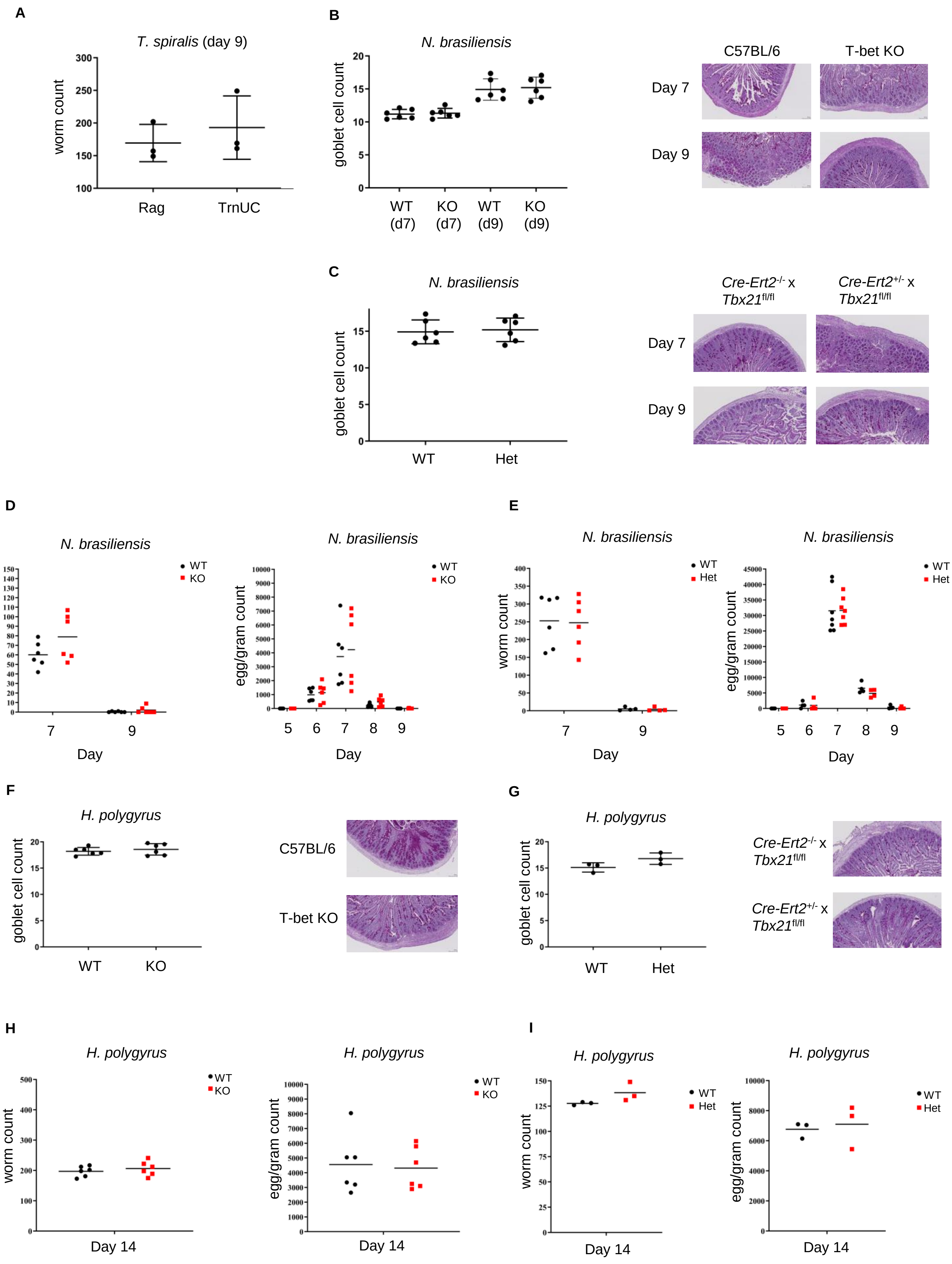

#### Supplementary Figure 3

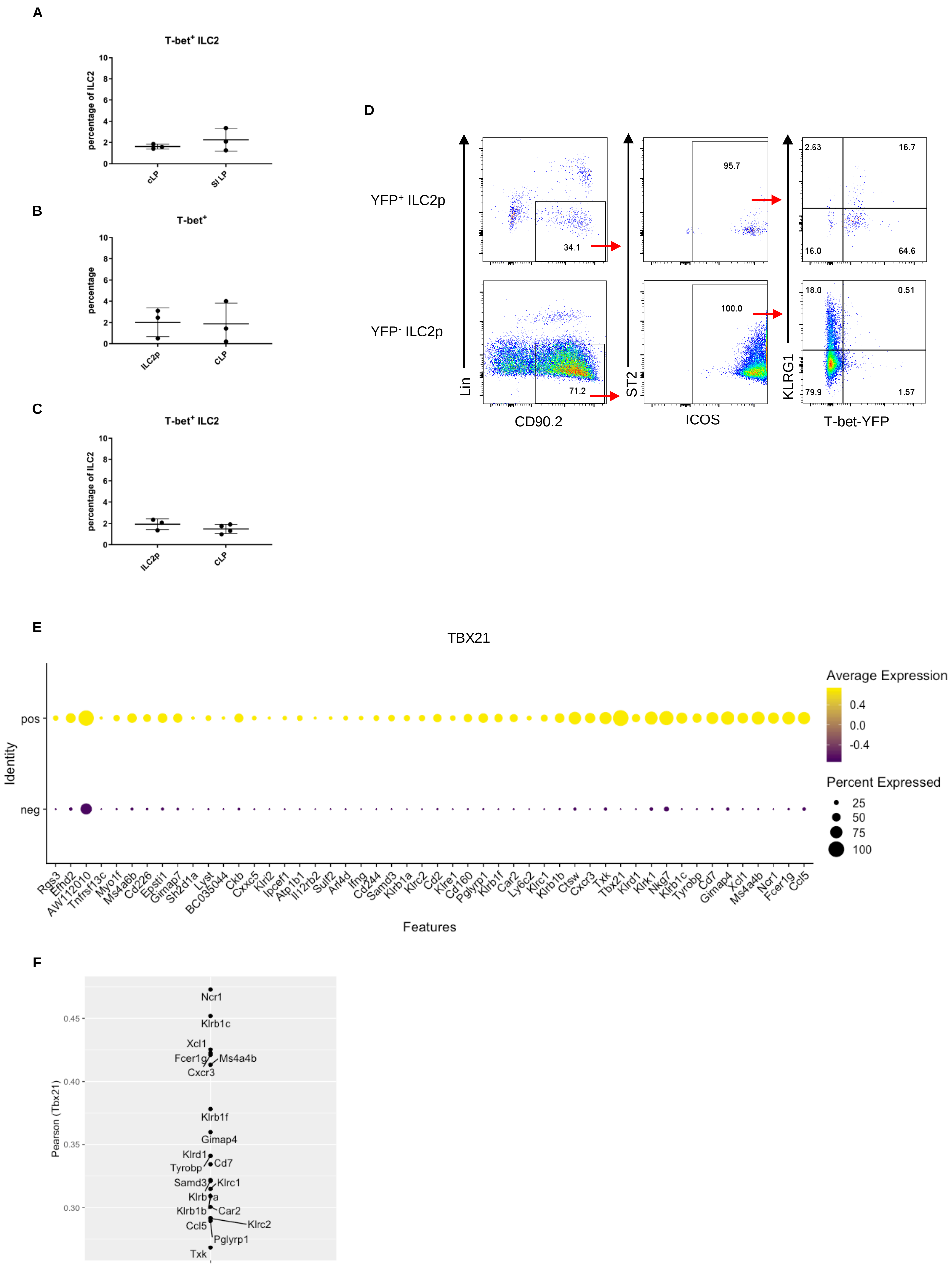
